# Characterizing upper extremity motor behavior in the first week after stroke

**DOI:** 10.1101/735431

**Authors:** Jessica Barth, Shashwati Geed, Abigail Mitchell, Peter S. Lum, Dorothy F. Edwards, Alexander W. Dromerick

**Author notes:** Corresponding author: (AWD).

## Abstract

**Background:** Animal models of brain recovery identify the first days after lesioning as a time of great flux in sensorimotor function and physiology; these findings have implications for human stroke recovery. After rodent motor system lesioning, daily skill training in the less affected forelimb reduces skill acquisition in the more affected forelimb. We asked whether spontaneous human motor behaviors of the less affected upper extremity (UE) early after stroke resemble the animal training model, with the potential to suppress clinical recovery.

**Methods:** This prospective observational study used a convenience sample of patients <7 days after stroke (n=25) with a wide severity range; Controls were hospitalized for non-neurological conditions (n=12). Outcome measures were Accelerometry, Upper-Extremity Fugl-Meyer (UEFM), Action Research Arm Test (ARAT), Shoulder Abduction/ Finger Extension Test (SAFE), NIH Stroke Scale (NIHSS).

**Results:** Accelerometry indicated total paretic UE movement was reduced compared to controls, primarily due to a 44% reduction of bilateral UE use. Unilateral paretic movement was unchanged. Movement shifted to unilateral use of the nonparetic UE, which increased by 77%. Low correlations between movement time and motor performance prompted an exploratory factor analysis (EFA) revealing a 2-component solution; motor performance tests load on one component (motor performance) whereas accelerometry-derived variables load on a second non-orthogonal component (quantity of movement).

**Conclusions:** Early after stroke, spontaneous overall UE movement is reduced, and movement shifts to unilateral use of the nonparetic UE. Thus, spontaneously-arising UE motor behaviors early after stroke are potential substrates for two mechanisms associated with poorer motor outcomes in animal models: learned non-use and inhibition of motor recovery through training of the nonparetic side. Accurate UE motor assessment requires two independent constructs: motor performance and quantity of movement. These findings provide opportunities and measurement methods for studies to develop new behaviorally-based stroke recovery treatments that begin early after onset.

## Introduction

Of the estimated 7 million stroke survivors in the United States, up to 88% are thought to have upper extremity (UE) motor involvement. This motor impairment is usually disabling, leading to the need for modification or assistance in activities of daily living (ADL) and reduced social participation (1-3) (4-6).

A series of animal studies have identified the first several days after stroke as a time of great flux in sensorimotor function and physiology; these processes have long term implications for recovery in animal models. A molecular program of injury and reparative responses has been well documented (7-13). Several motor behavioral factors (timing of motor training, dosing, and use of non-paretic forelimb) and various medications are active in meaningfully influencing forelimb motor recovery(14). These factors also seem relevant to conventional rehabilitation in clinical settings, with the potential to influence final motor outcomes in patients. One influential study examining the amount and timing of structured motor activity suggests that initiation of motor training beyond 5-14 days may be ineffective in restoring motor function, positing the existence of a critical period in stroke recovery(15). In contrast, other work found that overtraining early after stroke could actually lead to worse motor outcomes, and even activity-dependent lesion enlargement (16, 17). These study results are of course not mutually exclusive, and their relevance to humans with stroke remains unclear. Twenty minutes daily of skill training in the less affected forelimb at has been shown to reduce skill acquisition in the affected limb following a lesion of the motor cortex in rodents (18, 19). Were this true in human clinical populations, training compensation skills (in formal therapy or informally by a patient or family) in the nonparetic limb might inhibit motor recovery. The effects of early “nonuse” behaviors have been well established in non-human primate models (20-23).

In clinical populations, the data are limited and inconclusive. In the case of early initiation of rehabilitation, numerous retrospective studies have found that early rehabilitation admission is associated with better outcomes (9, 24-26). However, these mostly retrospective studies are confounded by stroke severity and medical complexity, since individuals who undergo rehabilitation sooner tend to be less severely affected and more medically stable. Formal randomized controlled trials involving dosing of rehabilitation during the first few days after stroke onset are limited but have been less positive. The Phase III A Very Early Rehabilitation Trial (AVERT) trial, in which patients were randomized to conventional and high intensity mobilization, found worse outcomes and greater mortality at 90 days for patients randomized to the high intensity group (27). Similarly, the Phase II Very Early Constraint-Inducted Movement During Stroke Rehabilitation (VECTORS) trial, in which patients were randomized at a mean of 9 ± 4.5 days after stroke, found that higher doses of motor training were associated with worse motor outcomes at 90 days (28, 29). Yet, other studies found no such effect (30).

Given the potential for these motor behaviors and interventions to affect motor recovery, systematic study of the human motor system in the first days after stroke is essential. In order to lay the groundwork for human trials, we undertook a preliminary study to identify and characterize the range of UE motor behaviors during the first week after stroke, using widely accepted clinical measures and also objective quantification using wrist-worn accelerometers. We focused on persons in the range of UE motor severity likely to need or receive rehabilitation interventions. We hypothesized that measures of motor impairment would be highly correlated, and that the amount of spontaneous movement would be both considerable and tightly linked to motor severity.

## Methods

### Subjects

The study was approved by the MedStar Health IRB. In this preliminary work, we enrolled a convenience sample with a wide variety of stroke severity, capturing a range of motor behaviors. Study participants were identified via screening logs maintained by the Stroke Central Atlantic Network for Research(31). A total of 448 patients were screened from MedStar Washington Hospital Center from April 16, 2017 to May 11, 2018; participants were consented ≤ 7 days of onset from the acute stroke service. All participants had imaging-confirmed unilateral ischemic or hemorrhagic stroke. Participants were excluded if they had a history of prior stroke with residual UE weakness, prior relevant orthopedic or neurological conditions that limited or potentially altered UE movement, and enrollment in a conflicting clinical trial. To control for non-specific UE use in hospitalized individuals, we recruited a second cohort of adult inpatients (*n*=12) with non-neurological conditions.

### Measures

*The Upper Extremity Fugl-Meyer* (UEFM) assessed motor impairment at the shoulder, elbow, wrist and fingers along with passive range of motion (PROM), pain and sensation. A full score is 66 with higher scores reflecting more normal motor function (32, 33).

*The Action Research Arm Test* (ARAT) assessed UE functional limitation. The ARAT uses a 4-point ordinal scale on 19 items to measure grasp, grip, pinch and gross motor movements of both UE(34, 35).

*The Shoulder Abduction-Finger Extension* (SAFE) strength assessment sums manual muscle testing of shoulder abduction and finger extension to produce a score from 0-10(36). A score of 10 indicates full strength in both movements.

*Goniometry* was performed according to standard methods at the wrist, fingers and elbow. (37).

*The National Institutes of Health Stroke Scale* (NIHSS), a standardized neurological assessment, described overall stroke severity. The UE motor item scores the participant on their ability to maintain shoulder flexion at 90 degrees and full antigravity extension of other UE joints for 5 seconds. Scores range from 0 (no impairment) to 4 (no movement).(38)

The *Mesulam Symbol Cancellation Test* (unstructured condition) screens for visuospatial neglect(39). This measure involves cancelling visual targets within a page of distractors; asymmetry of >3 errors indicates neglect.

*The Frenchay Aphasia Screening Test* (FAST) is a brief screening test for aphasia after stroke(40). The full scale is composed of four subscales; a higher score indicates more language impairment.

*The Wong-Baker Pain Rating Scale* was used to assess pain. (41, 42) *Accelerometry* (Actigraph GT9X Link Activity Monitor; Actigraph, Pensacola, FL) was used as an objective and quantitative measure of UE use bilaterally. Accelerations were recorded along 3 axes at 50Hz.

### Procedures

All participants or their proxy provided written informed consent and received standard acute stroke clinical care. A licensed Occupational Therapist performed all clinical assessments. Accelerometers were applied bilaterally just above the ulnar styli for at least 24 hours; data were normalized to 24 hours’ wear.

### Analysis

Immediately after removal of accelerometers from participants, data were downloaded and inspected visually for integrity using ActiLife software v6.13.3 (Actigraph, Pensacola, FL). ActiLife software converted the sample into 1Hz “counts” (one second epochs) for further analysis on subsequent programs. Counts represent the magnitude of acceleration after filtering to attenuate frequencies not associated with human movement. Data were then exported from the ActiLife software into a custom MATLAB program (Mathworks; Natick, MA) to calculate several previously reported metrics(43). The thresholding method reported by Urban and Lang(43) was used to determine if movement was present in each 1 second epoch. Total duration of movement of each limb was calculated, as well as duration of unilateral and bilateral movements. Separation of movement into unilateral only or simultaneous bilateral was possible since the data streams from the two accelerometers were synchronized. We calculated use ratios (activity ratio of hours of movement of the paretic limb to the non-paretic limb) to normalize variation in overall activity levels across subjects.

Descriptive statistics on participant demographics, clinical function tests, and accelerometry were calculated. Pearson’s correlations were computed between clinical function tests (UEFM, ARAT, and SAFE score) and the accelerometry measures of movement count and use ratio on the paretic side in the stroke group to determine strength of relationship between these measures. A Bonferroni correction was applied for multiple pairwise comparisons (adjusted *p* = 0.01, for five pairwise comparisons). Correlations showed non-significant relationships between the clinical function tests and accelerometry measures, prompting an unplanned, principal component factor analysis on these measures to determine if they measured similar or different constructs. The number of components to extract was decided using Kaiser’s Eigenvalue greater than 1 criteria (44, 45). A varimax orthogonal rotation was applied to improve interpretability of extracted factors. Factor loadings >0.4 were considered significant (46). All analyses were completed using IBM SPSS Statistics, Version 25.

## Results

Twenty-five participants were recruited at 4.5 ± 1.8 days after stroke onset (see Table 1). Stroke severity ranged from very mild to moderately severe (total NIHSS 0-19), and arm motor impairment was similarly distributed with UEFM scores ranging from 0-65. Of the 25 subjects, 8 had neglect and 8 reported mild to moderate pain. For controls, we recruited twelve acute rehabilitation inpatients (4 orthopedic, 5 leg amputation, 2 general debility and 1 cardiac). They had no neurological conditions and normal UE strength, sensation, and ROM on occupational therapy evaluation.

**Table 1:** Participant Characteristics.

| Demography | Side affected by stroke |  |  |  |
| --- | --- | --- | --- | --- |
|  | Controls<br>*n* = 12 | Stroke<br>*n* = 25 | Dominant<br>*n* = 10 | Non-dominant<br>*n* = 15 |
| Age | 61 (12.7)<br>33-85 | 58 (13.4)<br>33-85 | 62.5 (13.3)<br>48-85 | 57.8 (15.1)<br>33-83 |
| % Female | 66.6 | 36 | 50 | 26.7 |
| Days after onset | 7.6 (2.1)<br>6-13 | 4.5 (1.8)<br>1-7 | 4.7 (2.0)<br>1-7 | 4.5 (1.7)<br>1-7 |
| Motor performance measures |  |  |  |  |
| ARAT | - | 27.8 (22.6)<br>0-57 | 28.3<br>0-57 | 27.5 (23.7)<br>0-57 |
| UEFM | - | 35.4 (22.2)<br>0-65 | 34.4 (22.6)<br>0-65 | 36.1 (22.8)<br>6-64 |
| SAFE | 9.5 (.52)<br>9-10 | 5.7 (3.5)<br>0-10 | 5.7 (3.5)<br>0-10 | 5.7 (3.7)<br>0-10 |
| Active Wrist extension, degrees | - | 22.8 (18.1)<br>0-58 | 22.4 (18.1)<br>0-48 | 23.1 (18.1)<br>0-58 |
| NIHSS Total | - | 6.2 (4.7)<br>0-19 | 6.5 (5.5)<br>0-19 | 6.0 (4.3)<br>0-16 |
| NIHSS Motor Arm | - | 1.5 (1.7)<br>0-4 | 1.4 (1.3)<br>0-4 | 1.6 (4.3)<br>0-4 |

Demographics of study participants. Values are mean (SD) with ranges underneath.
Hand dominance was confirmed with patient or family. In the Control group, dominant hand served as nonparetic limb.

The amount of spontaneous UE movement as measured by accelerometry is presented in Table 2. We used the non-dominant UE activity of the control group as a benchmark (23, 47-51) for paretic limb activity in the stroke sample. Overall, stroke participants showed considerable movement (3.7 ± 3.1 hours) that was nonetheless significantly reduced on the paretic side compared to controls’ non-dominant side (6.2 ± 2.8 hours; *t* (35) = −2.29, *p* = 0.03). There was no significant difference between the amount of nonparetic UE movement in the stroke group compared to the dominant limb of controls (t (35) = −0.10, *p* = 0.92). This amount of UE movement is far less than community-based samples previously reported by us and others, which typically range from 3-6 hours(47, 52-54).

**Table 2:** 24 Hour Accelerometry Results. Units are in Hours of Movement per 24 hours.

|  | Control† | Acute<br>Stroke Pt<br>N=25<br>Mean (SD) | Dominant<br>Affected<br>n=10 | Non-Dominant<br>Affected<br>n=15 | Comparison<br>p-value |
| --- | --- | --- | --- | --- | --- |
| Total UE movement, paretic | 6.2 ± 2.8 | 3.7 ± 3.1 | 4.4 ± 3.9 | 3.0 ± 2.3 | 0.03* |
| Total# UE movement, non-paretic | 7.0 ± 2.5 | 6.9 ± 4.5 | 8.2 ± 5.7 | 6.1 ± 3.3 | 0.92 |
| Simultaneous Bilateral Movement | 5.0 ± 2.5 | 2.8 ± 2.5 | 3.7 ± 3.3 | 2.2 ± 1.7 | 0.02* |
| Paretic Unilateral | 1.1 ± .60 | 0.93 ± .90 | 1.0 ± .79 | .9 ± 1.0 | 0.41 |
| Non-Paretic Unilateral | 2.2 ± 1.2 | 3.9 ± 2.7 | 4.3 ± 3.4 | 3.7 ± 2.2 | 0.008** |
| Use Ratio | .89 ± .20 | .56 ± .30 | .58 ± .30 | .56 ± .37 | 0.001** |

†In Control subjects, the non-dominant limb was treated as the "paretic limb."

Next, we asked whether patients behaved in a way that could mimic the rodent model(18, 19) by increasing the unilateral use of their nonparetic limb and thus potentially engaging in skill training. Specifically, we examined whether patients compensated for hemiparesis by increasing bilateral UE movements or by increasing unilateral movements of the nonparetic UE. Comparison of unilateral use of the nonparetic UE in stroke patients (3.9 ± 2.7 hours) and dominant UE in controls (2.2 ± 1.2 hours) showed a significant increase in the amount of movement in the nonparetic UE in stroke patients (*t* (34.7) = 2.81, *p* = 0.008. Thus, stroke patients increased the unilateral use of their nonparetic limb (3.9 ± 2.7 hours) compared to controls (2.2 ± 1.2 hours). Moreover, there was a significant reduction of bilateral UE use in stroke (2.8 ± 2.5 hours) compared to controls (5.0 ± 2.5 hours; (*t* (21.98) = −2.532, *p* = 0.02. The use ratio in stroke participants (0.56 ± 0.34) was significantly lower compared to the control participants (0.89 ± 0.18, *t* [34.3] = −3.79, *p* = 0.001). Taken together these data indicate increased unilateral use of the nonparetic UE for compensation.

Next, we examined the relationship between UE motor function measured with performance scales and the amount of spontaneous movement measured with accelerometry. The correlation matrix (Table 3) shows strong significant correlations between UEFM, ARAT, SAFE, and NIHSS motor arm item (Bonferroni adjusted *p* = 0.01). Use ratio was moderately correlated with each of the performance scales. Notably, there was no significant difference in the paretic arm hours of movement (t-1.54 (22), p=0.14) in those with a high or low UEFM scores determined using a median split (high ≥33, low ≤32). Thus, in our sample assessed early after stroke, individuals with low UEFM scores moved their UE’s just as much as those with high UEFM scores who are often predicted to have better long-term recovery(47, 51, 55). This dissociation between performance scales and amount of spontaneous movement was an unexpected finding that suggested that these might measure different dimensions of UE impairment. To test this hypothesis, we computed a principal component factor analysis using an orthogonal (Varimax) rotation to UEFM, ARAT, SAFE scores, range of motion at wrist, range of motion at metacarpo-phalangeal joint, and hours of movement from paretic and non-paretic side, see Table 4.

**Table 3:** Correlations among Measures. Correlation matrix between clinical function tests and accelerometry-derived measures in Stroke group.

|  | ARAT | UEFM | SAFE | NIHSS<br>Motor arm | Use ratio | Hours of<br>Use of<br>Paretic<br>Limb |
| --- | --- | --- | --- | --- | --- | --- |
| ARAT | 1 |  |  |  |  |  |
| UEFM | 0.97** | 1 |  |  |  |  |
| SAFE | 0.88** | 0.93** | 1 |  |  |  |
| NIHSS Motor arm | - 0.85** | -0.87** | -0.89** | 1 |  |  |
| Use ratio | 0.65** | 0.69** | -0.69** | -0.69** | 1 |  |
| Hours of Use of<br>Paretic Limb | 0.32 | 0.34 | 0.49 | -0.48 | 0.51** | 1 |

\*\*Correlation is significant at 0.01 level, adjusted $p$ for multiple comparisons (5 pairwise comparisons, original $p = 0.05$ )

**Table 4:**
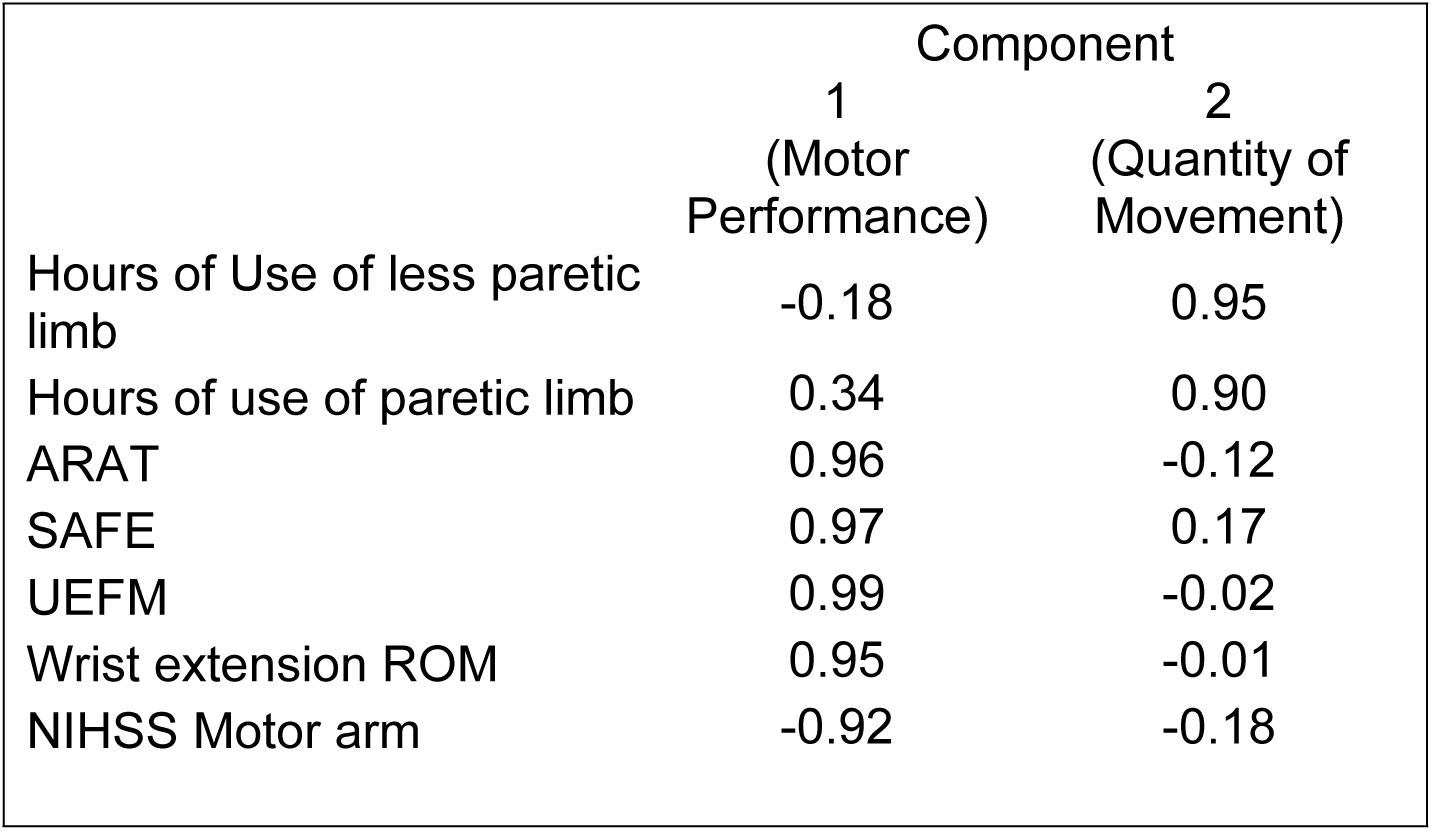
Item loadings based on two-factor solution with oblique rotation. Components 1 and 2 explain 92.8% of the variability in UE impairment measured by the combination of variables entered into the factor analysis. Hours of use of paretic and nonparetic sides load on Component 2 (“Quantity of movement”), suggesting these measures represent a distinct construct compared to variables loading on Component 1 (“Motor performance”)

The final rotated solution achieved a simple structure and revealed a two-factor solution, explaining 92.8% of the variability in UE impairment. Component 1 showed high factor loadings from UEFM, ARAT, SAFE, and NIHSS scales with range of movement (wrist extension). Given the predominance of variables assessing movement performance on this component, we labeled Component 1 as aligning with the latent construct of “Motor performance”. Hours of movement of the paretic and non-paretic UE load heavily on Component 2. Given that both these accelerometry measures align with the amount of movement, we labeled Component 2 as aligning with the latent construct of “Quantity of movement”. Overall, 92.8% of the variance of UE movement in the first week after stroke was explained by the sum of Motor performance factor (67.4%) and by the Quantity of movement (25.4%).

## Discussion

Animal models of stroke recovery demonstrate patterns of gene expression and injury responses in the first few days after injury that evolve with time(8, 13, 56, 57). These processes might constitute a time-limited “sensitive period” for enhanced recovery after stroke(26). In this pilot study of motor behaviors during the first week after stroke, we sought to characterize in patients the aspects of spontaneous motor behaviors (58, 59) that animal models suggest (18) influence motor outcome. Motor activity during this time window is poorly understood, perhaps because it falls outside the early intervention window of interest to vascular neurologists and before the substantive interventions of interest to rehabilitation investigators. We found several notable features.

First, motor performance measures are strongly correlated at this early post-stroke time-point, just as they have been demonstrated to correlate at later times after stroke(47, 60-62). This result will simplify future studies during this time period, allowing a lean UE assessment with fewer motor performance measures, reducing subject burden.

Second, we found patterns of compensation that raise the possibility of spontaneous behaviors that influence recovery in the affected limb. Specifically, movement shifted to the non-paretic limb, which was in motion 4.2 times more than the paretic side. This finding shows that even this early, two possible mechanisms that could influence motor recovery are already in place: 1. the occurrence of learned non-use in the affected UE and 2. skill acquisition in the non-paretic limb that could impede recovery. Further supporting the picture of non-paretic limb compensation, the amount of bilateral simultaneous movements was reduced compared to controls, suggesting that bilateral movements were not the preferred compensation strategy during this time period.

Finally, the factor analysis identified two latent constructs underlying measurement of UE impairment. We characterized Component 1 as aligning with “Motor performance’ and Component 2 as aligning with “Quantity of movement”. Motor performance, measured by a combination of clinical function tests like UEFM, ARAT, and SAFE accounts for 67% of the variance in UE impairment; whereas Quantity of movement, measured by accelerometry, accounts for an additional 25% of the variance. This two-factor solution is a parsimonious solution as evidenced by the amounts of variance explained by each factor. A third component only added an additional 2% variance in UE impairment compared, suggesting the 2-factor solution provided the optimal solution. The orthogonal rotation resulted in motor performance measures aligning to Component 1 and accelerometry measures aligning to Component 2. Although it is non-traditional for a “factor” to have only two variables with significant loadings as in the case of our Component 2, this two-factor solution is statistically and clinically meaningful. Together, the different metrics we used could account for approximately 93% of the variance observed in UE impairment within a week after stroke. These results highlight the need to measure both motor performance and quantity of movement when evaluating UE function during this post-stroke period. Future work should determine the relative importance of these constructs and determine whether other constructs not measured in this study may also be present.

Our work extends that of Gebruers et al. (63) who used different accelerometry analyses that make direct comparisons difficult. Our results are generally consistent with their correlations between use ratio and UEFM. However, we did not find a correlation between hours of movement and UEFM. The differences arise because we used Pearson’s correlation, given the non-parametric nature of UEFM. When we tested correlations between hours of movement and UEFM using Spearman’s test, our findings were consistent with Gebruers et al(55).

Our approach also explains why Gebruers et al.(63) found that hours of movement were not good predictors of disability at 3-months. Gebruers et al. found hours of motion were a good predictor only when there was going to be a “good outcome”, not if there was a “bad outcome” at 3 months. Our factor analysis explains this dissociation between the good outcome and bad outcome groups: hours of movement does not measure the same construct as UEFM. In individuals with high UEFM scores, hours of movement are highly correlated with performance measures. However, in participants with low UEFM scores, there is only a weak correlation with hours of movement, providing a poor prediction of outcome at 3 months. Our analysis emphasizes this dissociation; there is no significant difference between hours of movement in those with high versus low UEM scores. Thus, measuring both hours of movement and motor performance is required to realistically assess motor function. Use ratios are less affected, because they are loaded on the motor performance axis, similar to the UEFM.

Our results contribute to the systematic examination and application of specific animal model findings to the challenging clinical context of the first week post-stroke (26, 64). Animal data clearly indicate that the dose of formal motor training early after lesioning can affect motor outcome in a variety of ways depending on timing and dose of training, and which forelimb is trained(15, 17-19, 26). Non-paretic forelimb skill training has been shown to inhibit motor recovery of the paretic rodent forelimb; if these findings hold in humans, this compensation may potentially worsen long term UE motor outcome in the clinical setting. Our human data indicate reduced but substantial spontaneous UE motor activity with a shift to the non-paretic UE, providing the substrate for spontaneous unintended training of the non-paretic limb(18, 19). This behavior also indicates that the behavioral substrate for the learned nonuse phenomenon is present quite early after injury (22, 65, 66).

At this time, our data cannot predict whether these UE behaviors are therapeutic, deleterious, or simply irrelevant to long term motor outcomes. Further research is needed to discern whether in clinical populations there is an impact on eventual motor outcome. If so, development of clinical interventions could be required.

### Study Limitations

The sample recruited from an urban safety net hospital may not be representative of all stroke patients, and future studies should recruit a larger and more representative sample. The cross-sectional design and 24-hour assessment period do not allow assessment of the rapid motor changes seen in the first days of stroke. Our attempts to collect 30-day outcome data were hampered by participant unavailability, death or recurrent stroke, and lack of recall of study consent by some participants and families. Other constructs relevant to UE behavior may exist.

## Conclusion

With this study we were successful in recruiting and testing stroke subjects within one week of onset. The sample ranged from very mild to severely impaired participants as measured by highly correlated motor performance measures. Accelerometry metrics in these early days after stroke show an overall reduction of bilateral UE movement with individuals compensating by shifting their UE movement to the nonparetic limb.

Compensatory strategies chosen by patients provide the substrate for potential detrimental behaviors discovered in animal recovery models: learned non-use and inhibition of motor recovery by training of the unimpaired forelimb. Finally, factor analysis shows that assessment of motor performance alone is insufficient to describe UE movement, adding accelerometry explains 93% of the variance completing the assessment of UE behavior early after stroke.

## Data Availability

All relevant data are within the paper and its Supporting Information files. Raw data set from accelerometry recordings and demographics are available at figshare:

https://figshare.com/projects/Characterizing_upper_extremity_motor_behavior_in_the_first_week_after_stroke/63638

## Financial Disclosure Statement

The following work was funded by the following:

Center for Brain Plasticity and Recovery (JB, SG, AM, PL, DE, AD)

Stroke Central Atlantic Network for Research: NINDS NS086513 (JB, SG, DE, AD) Full names of commercial companies that funded the study or authors (N/A)

URLs to sponsors’ websites: https://cbpr.georgetown.edu/

https://www.ninds.nih.gov/

The funders had no role in study design, data collection and analysis, decision to publish or preparation of the manuscript.

